# Allele frequency differentiation at height-associated SNPs among continental human populations

**DOI:** 10.1101/2020.09.28.317552

**Authors:** Minhui Chen, Charleston W. K. Chiang

## Abstract

Polygenic adaptation is thought to be an important mechanism of phenotypic evolution in humans, although recent evidence of confounding due to residual stratification in consortium GWAS made studies of polygenic adaptation more difficult to interpret. Using F_ST_ as a measure of allele frequency differentiation, a previous study has shown that the mean F_ST_ among African, East Asian, and European populations is significantly higher at height-associated SNPs than that found at matched non-associated SNPs, suggesting that polygenic adaptation is one of the reasons for differences in human height among these continental populations. However, we showed here even though the height-associated SNPs were identified using only European ancestry individuals, the estimated effect sizes are significantly associated with structures across continental populations, potentially explaining the elevated level of differentiation previously reported. To alleviate concerns of biased ascertainment of SNPs, we re-examined the distribution of F_ST_ at height-associated alleles ascertained from two biobank level GWAS (UK Biobank, UKB, and Biobank Japan, BBJ). We showed that when compared to non-associated SNPs, height-associated SNPs remain significantly differentiated among African, East Asian, and European populations from both 1000 Genomes (*p* = 0.0012 and *p* = 0.0265 when height SNPs were ascertained from UKB and BBJ, respectively), and Human Genome Diversity Panels (*p* = 0.0225 for UKB and *p* = 0.0032 for BBJ analyses). In contrast to F_ST_-based analyses, we found no significant difference or consistent ranked order among continental populations in polygenic height scores constructed from SNPs ascertained from UKB and BBJ. In summary, our results suggest that, consistent with previous reports, height-associated SNPs are significantly differentiated in frequencies among continental populations after removing concerns of confounding by uncorrected stratification. Polygenic score-based analysis in this context appears to be susceptible to the choice of SNPs and, as we compared to F_ST_-based statistics in simulations, would lose power in detecting polygenic adaptation if there are independent converging selections in more than one population.

## Introduction

Because of the highly polygenic nature of many human complex traits, polygenic adaptation was thought to be an important mechanism of phenotypic evolution in humans. As each genetic locus contributes a small effect to complex traits, polygenic adaptation is expected to be different from the classical selective sweep in that only a subtle but coordinated allelic or haplotypic signature across loci underlying the selected trait is expected (Pritchard *et al*. 2010).

In humans, height is one of the earliest putative examples of polygenic adaptation (Turchin *et al*. 2012; Berg and Coop 2014; Robinson *et al*. 2015; Field *et al*. 2016; Guo *et al*. 2018). By evaluating allele frequency differentiations at height-associated SNPs, a previous study suggested polygenic adaptation as one of the reasons for differences in human height among global populations (Guo *et al*. 2018). Specifically, Guo et al. demonstrated that compared to randomly selected, frequency- and LD-score-matched SNPs, height-associated SNPs showed significantly higher mean F_ST_ across the three continental populations (Africans, Europeans, and East Asians). However, height-associated SNPs examined in Guo et al. were ascertained from consortium genome-wide association studies (GWAS) such as the Genetic Investigation of Anthropometric Traits (GIANT) (Wood *et al*. 2014). Previous studies have suggested that because of residual uncorrected stratification, the estimated effect sizes of SNPs detected in GIANT showed a subtle but biased correlation with population structure in Europe. As a result, polygenic scores (PSs) constructed on the basis of these SNPs showed exaggerated difference in human height between Northern and Southern Europeans (Berg *et al*. 2019; Sohail *et al*. 2019; Chen *et al*. 2020). Therefore, there is concern whether the previous polygenic signals among global populations were confounded by population stratification. On the one hand, while population stratification within Europe is not expected to bias allele frequency differentiations at height-associated SNPs among global populations, it is possible that structures among global populations are correlated with structures within Europe due to, for example, migration and admixture between Europeans and populations outside of Europe. This could result in spurious signals of adaptation. On the other hand, since F_ST_, the main statistics used by Guo et al. to measure allele frequency differentiations, is unsigned and does not rely on the estimated effect sizes, it could be more robust to residual stratifications in GWAS. It is important to note that the signals of adaptation were inferred at height-associated SNPs; height itself might not be the target of selection because it could be due to a trait that shares genetic architecture with height. Nevertheless, these inferred signals of adaptation suggest that natural selection contributed to the differentiation of height between human populations.

In the present study, we used summary statistics from UKB and BBJ to re-examine whether height-associated SNPs exhibit signs of adaptation among the three continental populations from Guo et al., i.e. Africans, Europeans and East Asians. We demonstrated that the effect sizes of height-associated SNPs ascertained from UKB and BBJ were much less associated with the population structure across the three continents, alleviating any concerns of confounding due to residual stratification. Using this approach, we found that allele frequencies of height-associated SNPs remain significantly differentiated among continental populations when compared to that of non-associated SNPs, consistent with a polygenic adaptive signal at these SNPs (Guo *et al*. 2018). However, applying the PS-based testing framework from either Guo et al. or Berg and Coop, we detected no significant difference in height PSs among the three populations and observed that even the ranked order of PSs among populations appeared sensitive to the choice of variants for analysis. This is consistent with the poor transferability of PSs across populations (Martin *et al*. 2017; but see Ragsdale *et al*. 2020), which could lead to a loss of power in PS-based framework. Alternatively, through simulations we also show that PS-based framework could lose power if different parts of the trait architecture is under selection in each of the populations undergoing independent convergent evolution. In this scenario F_ST_-based statistic would be more powered to detect evidence of adaptation.

## Materials and Methods

### GWAS Panels

To evaluate polygenic adaptation at height-associated SNPs, we obtained GWAS summary statistics from three studies. Briefly, they are:

1. GIANT (Wood *et al*. 2014), a meta-analysis of 79 separate GWASs for height using a total of ∼253,000 individuals of European ancestry with ∼2.5 M variants. Each study imputed their genetic data to HapMap Phase II CEU (Utah residents with ancestry from northern and western Europe) genotypes and then tested for association with sex-standardized height, assuming an additive inheritance model and adjusting for age and other study-specific covariates (including principal components [PCs]). Summary statistics were downloaded on 7/25/2019 from https://portals.broadinstitute.org/collaboration/giant/images/0/01/GIANT_HEIGHT_Wood_et_al_2014_publicrelease_HapMapCeuFreq.txt.gz.
2. UKB (http://www.nealelab.is/blog/2017/7/19/rapid-gwas-of-thousands-of-phenotypes-for-337000-samples-in-the-uk-biobank), a GWAS based on ∼337,000 individuals of white British ancestry in the UK Biobank. Genotyped individuals were imputed to whole-genome sequencing data from Haplotype Reference Consortium (HRC), UK10K (coverage = 7×), and 1000 Genomes for ∼10.8 M variants. Association testing was done on standardized height correcting for sex and 10 PCs. Summary statistics were downloaded on 7/15/2019 from https://www.dropbox.com/s/sbfgb6qd5i4cxku/50.assoc.tsv.gz.
3. BBJ (Akiyama *et al*. 2019), a GWAS based on ∼159,000 individuals of Japanese ancestry from Biobank Japan (Nagai *et al*. 2017; Hirata *et al*. 2017). Genotyped individuals were imputed to combined whole-genome sequencing data from BBJ1K (coverage = 30×) (Okada *et al*. 2018) and 1000 Genomes for ∼27.9 M variants. Individuals not of Japanese origins were excluded by self-report or principal component analysis (PCA). Using standardized residuals of height after adjusting for age, age^2^, and sex, a GWAS was conducted using a linear mixed model implemented in the software BOLT-LMM to control for cryptic relatedness and population structure (Loh *et al*. 2015). Summary statistics were downloaded on 6/19/2020 from https://humandbs.biosciencedbc.jp/files/hum0014/hum0014.v15.ht.v1.zip.

### Population genetic data

To evaluate polygenic selection at height-associated SNPs across continental populations, we analyzed samples from Africa, East Asia, and Europe from 1000 Genomes phase 3 release (1000 Genomes Project Consortium *et al*. 2015), including 504 individuals from five African subpopulations: ESN (Esan in Nigeria), GWD (Gambian in Western Divisions in the Gambia), LWK (Luhya in Webuye, Kenya), MSL (Mende in Sierra Leone), and YRI (Yoruba in Ibadan, Nigeria); 404 individuals from four European subpopulations: CEU, GBR (British in England and Scotland), IBS (Iberian Population in Spain), and TSI (Toscani in Italia); and 504 individuals from five East Asian subpopulations: CDX (Chinese Dai in Xishuangbanna, China), CHB (Han Chinese in Beijing, China), CHS (Southern Han Chinese), JPT (Japanese in Tokyo, Japan), and KHV (Kinh in Ho Chi Minh City, Vietnam). We did not include the two admixed African subpopulations (i.e. ACB [African Caribbeans in Barbados] and ASW [Americans of African Ancestry in SW USA], see **Figure S1**) and the FIN (Finnish in Finland) subpopulation because of its known unique demographic history (Wang *et al*. 2014; Locke *et al*. 2019). To confirm the signals of polygenic adaptation, we replicated the analysis on the Human Genome Diversity Project (HGDP) dataset (Bergström *et al*. 2020), using either all 929 samples from seven regions (including 104 Africans, 61 Americans, 197 Central South Asians, 223 East Asians, 155 Europeans, 161 Middle East, and 28 Oceanians), or 482 samples from three continents (104 Africans, 223 East Asians, and 155 Europeans).

### Population structure analysis

We first conducted PCA on the three continental populations (Europeans, Africans, and East Asians) from 1000 Genomes. We used biallelic SNPs with MAF > 5% in the dataset, and thinned to no more than one SNP in 200 kb. We did not prune the dataset by LD due to the mixture of continental populations. We further removed SNPs in known regions of long-range LD (Price *et al*. 2008). PCA was performed on the remaining variants via Eigensoft (version 7.2.1). We also conducted two within-continent PCAs in the same manner, one based on the four European subpopulations (CEU, GBR, IBS, and TSI) and the other based on the five East Asian subpopulations (CDX, CHB, CHS, JPT, and KHV) from 1000 Genomes. For the two within-continent PCAs, pruning the dataset by LD (using --indep-pairwise 50 5 0.2 in PLINK (Chang *et al*. 2015)) instead of by distance of 200 kb did not qualitatively change the results (results not shown).

A measure of potential confounding due to residual stratification is the correlation between estimated SNP effect sizes from GWAS summary statistics with its loading on a particular PC. A strongly significant correlation implies a systematic bias in effect sizes that aligns with the structure captured by a particular PC, potentially confounding the inference of polygenic adaptation (Sohail *et al*. 2019; Chen *et al*. 2020). To compute the correlation between PC loadings and SNP effect sizes from each GWAS panel, we first performed linear regressions of the PC value on the allelic genotype count for each variant polymorphic in the PCA populations, and we used the resulting regression coefficients as the variant’s PC loading estimates on a particular PC. For each PC and each GWAS panel, we then computed the Pearson correlation coefficient of PC loading and effect sizes, and tested whether the correlation coefficient is significantly different from zero on the basis of jackknife standard errors computed by splitting the genome into 1,000 blocks with an equal number of variants.

### Signature of Selection at Height-Associated SNPs

To ascertain height-associated variants, we selected a set of genome-wide significant variants (*p* < 5e-8) with MAF > 1% in GWAS panel after greedily pruning any other variants such that no two variants were within 1 Mb of each other. We then further pruned by LD using 1000 Genomes as the reference such that no two variants would have a r^2^ > 0.1. We used CEU, GBR, and JPT data as reference LD for pruning GIANT, UKB, and BBJ summary statistics, respectively. In total, we identified 458, 709, and 407 independent height-associated SNPs from GIANT, UKB, and BBJ summary statistics. To evaluate the evidence of selection at height-associated SNPs, we applied the following two approaches: F_ST_ enrichment test and PS-based tests.

#### F_ST_ enrichment test

We used Wright’s fixation index (F_ST_) calculated from PLINK (Chang *et al*. 2015) to measure the extent to which a particular SNP varied in allele frequency among the three continental populations from 1000 Genomes. Following Guo et al. (Guo *et al*. 2018), we compared the mean F_ST_ value of the height-associated SNPs with that of the presumed-neutral, non-associated, SNPs matched by MAF and LD score. First, we divided all the SNPs into 500 bins (25 bins based on MAF in increments of 0.02, but excluding SNPs with MAF < 0.01; within each MAF bin, there are 20 bins of LD scores in increments of 5% quantiles of LD score distribution). MAFs were obtained from GWAS summary statistics, and the LD scores were computed from 1000 Genomes as the sum of the LD *r*^*2*^ between the focal SNP and all the flanking SNPs within 10-Mb window. We used EUR (excluding FIN) as reference LD for GIANT and UKB, and used EAS for BBJ. Second, we allocated the height-associated SNPs to the MAF and LD stratified bins, randomly sampled a matched number of SNPs from each bin to compute an overall mean F_ST_ value for the “neutral” SNP set. This process was iterated 10,000 times to generate a distribution of mean F_ST_ under neutrality. Finally, a P-value was computed from a two-tailed test by comparing the observed mean F_ST_ value for the height-associated SNPs against the null distribution. We also conducted the same test among the three continental populations and among the seven regional populations from HGDP, separately.

#### PS-based analysis

We estimated the PS for each population as the sum of allele frequencies at a set of *L* height-associated SNPs weighted by effect sizes from each GWAS panel (i.e., 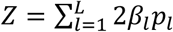, where *p*_*l*_ and *β*_*l*_ were the allele frequency and effect size at SNP *l*). We conducted the Q_X_ test as previously described (Berg and Coop 2014) to determine whether the estimated PSs exhibited more variance among populations than null expectation under genetic drift. The Q_X_ statistics follows a *χ*^2^ distribution with *M-1* degrees of freedom under neutrality, where *M* is the number of test populations, from which an asymptotic *p* value was estimated. Significant excess of variance among populations would be consistent with the differential action of natural selection among populations. To identify outlier populations and regions which contributed to the excess of variance, we further estimated the conditional *Z*-score proposed by Berg and Coop (Berg and Coop 2014). An extreme *Z*-score would suggest that the excluded populations had experienced directional selection on the trait of interest that was not experienced by the conditioned populations in the analysis. The scripts we used to implement these analyses are available on GitHub (see https://github.com/jjberg2/PolygenicAdaptationCode).

We also adopted the PS-based method used in Guo et al. (Guo *et al*. 2018) to estimate the deviation of the PS of a population from the overall mean. We calculated the PS for each individual as 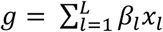 (where *L* is the number of height-associated SNPs, *x* represents the allele count at the SNP (0, 1, or2) and *β* is the estimate of SNP effect from the GWAS summary data), standardized the PS, and computed the mean PS for each population. To estimate whether an observed mean PS was significantly different from the null expectation under genetic drift, we randomly sampled a matched number of presumed neutral SNPs with the same strategy used in F_ST_ enrichment analysis, and calculated the mean PS for each population. We repeated this process 10,000 times to generate a distribution of the mean PS for each population. For each population, a *P*-value was computed from a two-tailed test by comparing the observed mean PS for the height-associated SNPs against the null distribution based on the neutral SNPs.

### Forward simulation

To evaluate the differences in power to detect polygenic adaptation under convergent evolution in multiple populations, we performed forward simulations in SLiM 3 (Haller and Messer 2019), assuming a three-populations model of constant *N*_*e*_ = 10^4^ that diverged 1,000 generations ago (**Figure S2**). We simulated 10,000 independent loci of 1Mb in length under neutrality with mutation rate = 2.36e-8 mutation/bp/generation, and recombination rate = 1e-8 recombination/bp/generation. We also simulated 120 independent loci of 100 kb in length under selection. We assumed the ancestral population was evolving neutrally prior to the population split 1,000 generations ago. At the split, we randomly chose one SNP with MAF > 20% from each of the 120 loci to be under selection. We simulated three scenarios involving different populations under selection. In the first scenario, all 120 SNPs were under selection in one population (P1). For half of the SNPs the derived alleles were beneficial and for the other half the derived alleles were deleterious. In the second scenario, two populations (P1 and P2) were under selection, each with a mutually exclusive set of 60 SNPs under selection (again half are beneficial, and half are deleterious). In the third scenario, three populations (P1, P2, and P3) were under selection, each with a mutually exclusive set of 40 SNPs under selection (20 beneficial and 20 deleterious mutations). We tested a range of selective coefficients (*s* = 1e-2, 5e-3, 2e-3, 1e-3, 8e-4, 4e-4, and 1e-4), assuming the same coefficient for every SNP under selection. We performed 100 independent replicates under each case. For each replicate, we sampled 500 individuals from each population, based on which we performed F_ST_ enrichment test and Q_x_ test on the 120 SNPs. We estimated their powers to detect signature of polygenic adaptation as the proportion of tests with positive signal (p < 0.05) under each scenario. In both tests we matched the selected alleles to separately simulated neutral alleles by MAF in P1, in effect assuming P1 is our “GWAS population” for ascertainment of trait-associated SNPs, though we had perfect power to detect all trait-associated SNPs. For the Qx test we also assumed that all beneficial and deleterious mutations had the same effect size of 1 and −1, and we further calculated the conditional *Z*-score in order to evaluate if conditional *Z*-score can correctly reflect outlier populations.

## Results

### Population stratification

The main conclusion of Guo et al. is that there is an association between the F_ST_ of a SNP among three continental populations and the magnitude of its effect size for height. That is, compared to matched SNPs from the rest of the genome, height-associated SNPs show significantly higher mean F_ST_ among the three continental populations. This conclusion may potentially be spurious, if there are significant correlation between effect sizes and the global population structure of the three continental populations. We have shown that effect sizes of height-associated SNPs ascertained from GIANT used by Guo et al. are significantly correlated with population structure within Europe (Sohail *et al*. 2019; Chen *et al*. 2020), but the correlation with global structure has not been examined.

We first evaluated the impact of population stratification on height-associated variants ascertained from different GWAS panels: the GIANT consortium, the UKB, and the BBJ datasets. We conducted PCAs across continents (Africa, Europe, and East Asia, **Figure S3**), within Europe (**Figure S4**), and within East Asia (**Figure S5**) using data from 1000 Genomes, and examined the correlation between effect sizes estimated from each GWAS panel and the PC loading on those PCAs. In the within-Europe analysis, GIANT effect sizes were significantly correlated with the loading of the first two PCs in Europe (rho = 0.124, p = 9.80e-92 for PC1; rho = 0.016, p = 1.70e-3 for PC2; **Figure S6**), as previously reported (Chen *et al*. 2020). We also observed that BBJ effect sizes were significantly correlated with the first two PCs in the analysis within East Asia (rho = −0.0182, p = 1.70e-5 for PC1; rho = −0.0288, p = 8.56e-10 for PC2; **Figure S7**), though the magnitudes of correlation were much lower compared to the correlation observed between GIANT effect sizes and PC1 in Europe (rho ∼ 0.02-0.03 vs. rho ∼ 0.12 in **Figure S6**). This could suggest that measures to control for population structure in BBJ were not completely effective in controlling for stratification within the East Asian continent, although the effect sizes estimated in BBJ are uncorrelated with population structure within Europe presumably due to its geographic distance (Chen *et al*. 2020).

In the analysis across continents, the first two PCs reflected the differentiation between Africans and Eurasians and the differentiation between Europeans and East Asians (**Figure S3**). We found that the effect sizes estimated in GIANT were highly correlated with the loading of the first two PCs of population structure (rho = 0.026, p = 1.13e-8 for PC1; rho = −0.086, p = 1.62e-78 for PC2; **Figure 1**), as well as with lower PCs that are driven by within-European structure (PC6 and PC8; **Figure 1 and S3**) or within-African structure (PC5; **Figure 1 and S3**). This suggests that even though consortium GWAS were conducted in European-ancestry populations, residual stratification can lead to a correlation between effect sizes of SNPs with inter-continental structures. Because F_ST_ measures are strongly associated with PC loadings (**Table S1**), the elevated differentiation among height-associated SNPs as reported by Guo et al. could potentially be a confounded observation.

**Figure 1.**
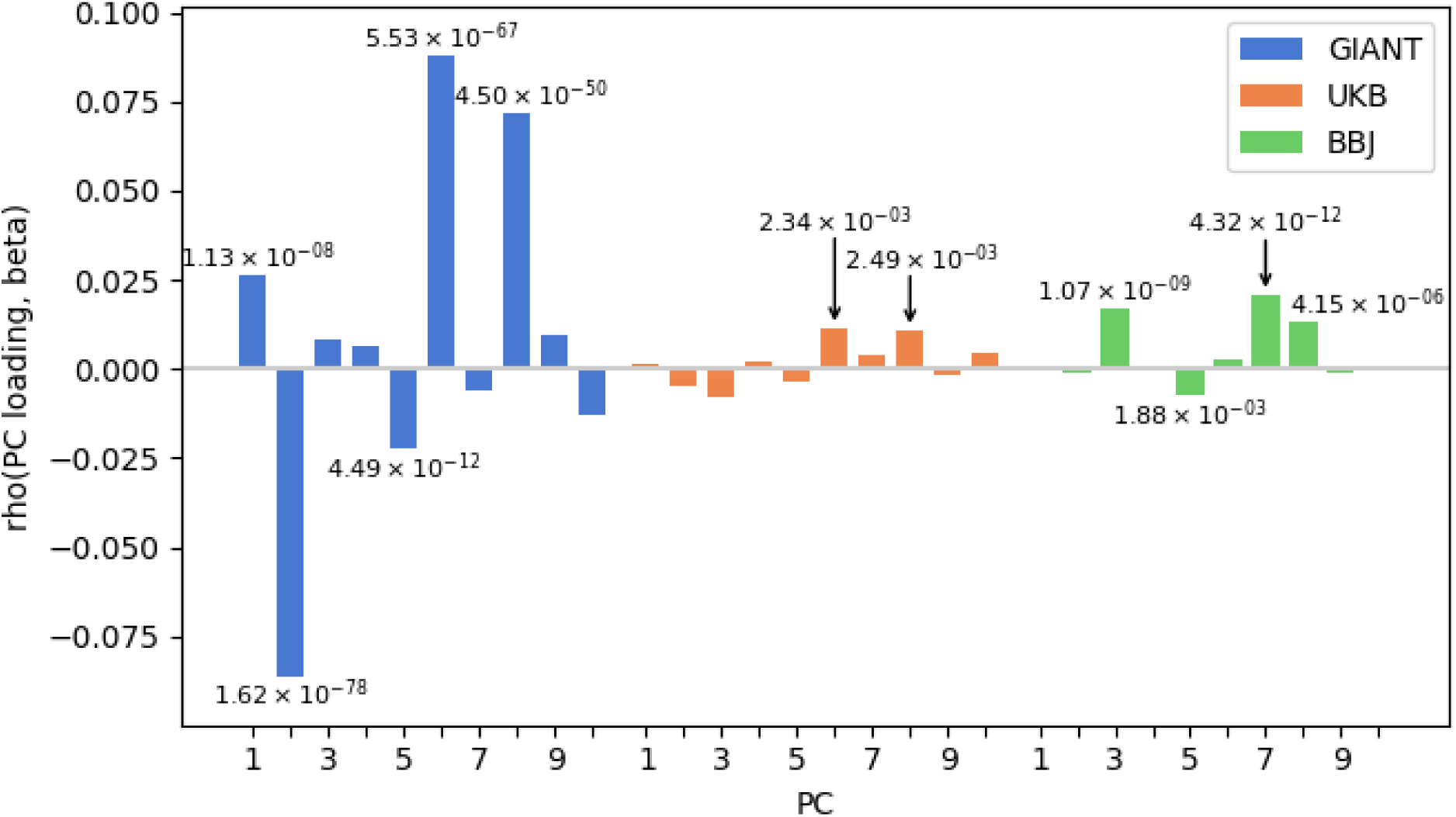
Evidence of stratification in GWAS summary statistics. Pearson correlation coefficients of PC loadings and SNP effects from GIANT, UKB, and BBJ based on all SNPs with MAF > 1% in each GWAS panel. Twenty PCs were computed in Africans, Europeans, and East Asians from 1000 Genomes; only the first 10 are shown here for readability. *P* values are based on jackknife standard errors (1,000 blocks). *P* values lower than 0.05/20 are indicated on each bar.

Compared to the situation in GIANT, the correlations were much smaller and insignificant in UKB (rho = 0.0016, p = 0.683 for PC1; rho = −0.0051, p = 0.107 for PC2) and BBJ (rho = −7.15e-4, p = 0.831 for PC1; rho = 0.0013, p = 0.673 for PC2). Even when there were significant associations between effect sizes and PC loadings of lower PCs, the magnitudes of the associations were much smaller (**Figure 1**). Therefore, UKB and BBJ summary statistics are not likely to be affected by population stratifications across continents and can be used to test the robustness of conclusions from Guo et al. based on F_ST_ differentiation.

### Enrichment of F_ST_ in height-associated SNPs

Because previous conclusions may have been spurious due to residual stratification, on the basis of independent SNPs associated with height with *p* < 5e-8 in UKB (709 SNPs) and BBJ (407 SNPs), we tested whether height-associated SNPs were more differentiated across continents than control SNPs, as measured by F_ST_. We found that the mean F_ST_ values for the height-associated SNPs ascertained from UKB and BBJ were significantly higher than those of the matched, presumed neutral, SNPs with *p* = 0.0012 in UKB and *p* = 0.0265 in BBJ (**Figure 2**), indicating that the SNPs associated with height were more differentiated. The signature using the height-associated SNPs ascertained from GIANT was weaker (*p* = 0.0621) compared to the signal (*p* = 4.93e-6) observed in the previous report (Guo *et al*. 2018). This is likely due to the previous authors using less stringent pruning parameter, less stringent *p*-value threshold, and including admixed 1000 Genomes populations in the previous analysis. This contributed to our analysis using ∼450 height-associated SNPs for analysis, compared to ∼1100 SNPs used in analysis in Guo et al., potentially decreasing our power. When we increased the *p* threshold for ascertaining height-associated SNPs to 5e-6 (681 SNPs), the same value was used in Guo et al. (Guo *et al*. 2018), the signal of F_ST_ enrichment test improved to *p* = 1.4e-3 (**Figure S8**). We also replicated our finding using data from HGDP. The mean F_ST_ of the height-associated SNPs remained significantly larger than that of control SNPs on the basis of summary statistics from UKB and BBJ with *p* = 0.0225 and *p* = 0.0032 for the analysis on the three continental populations and *p* = 0.0051 and *p* = 0.0002 for the analysis on the seven regional populations (**Figure 3**).

**Figure 2.**
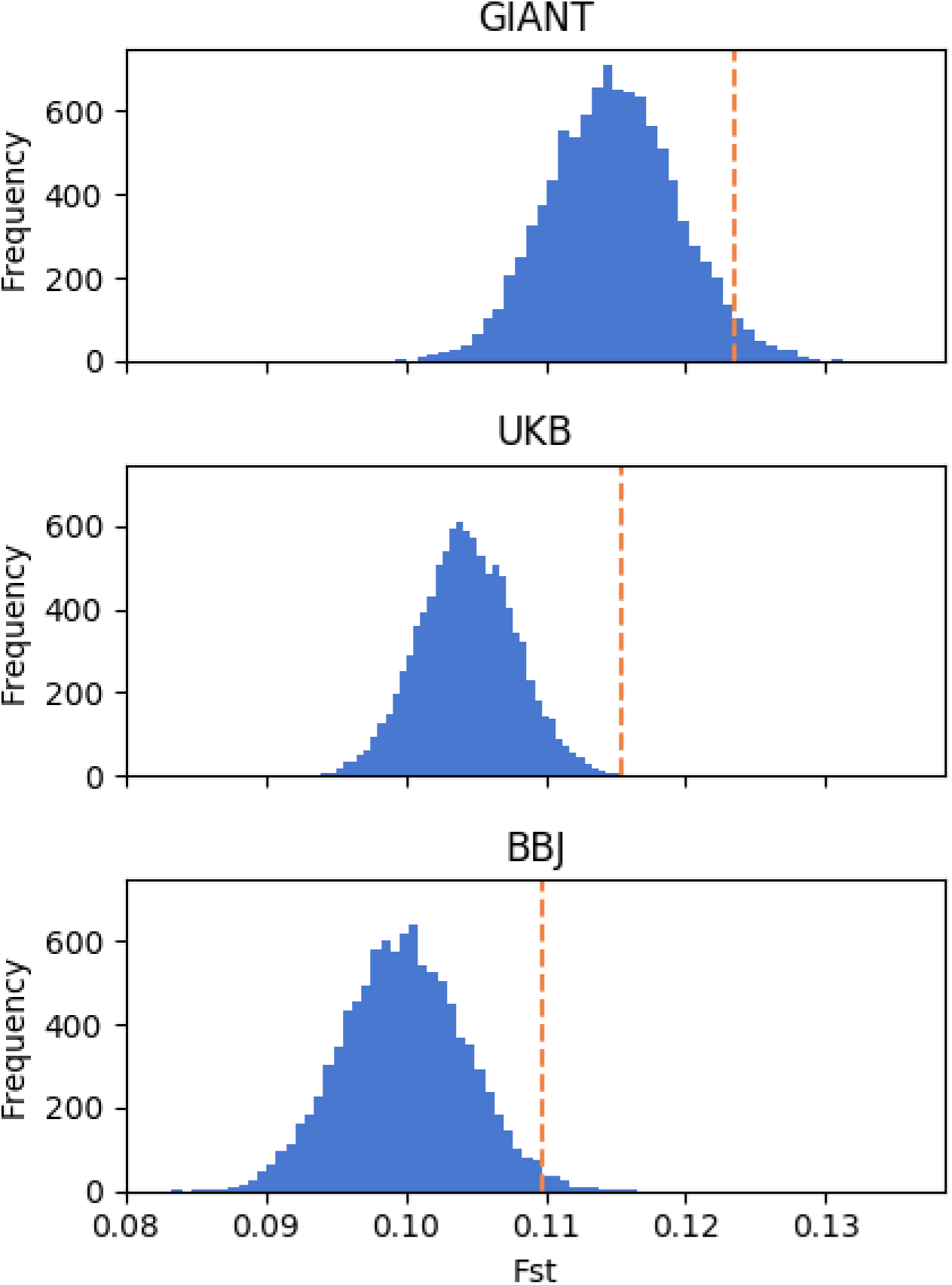
Mean F_ST_ values of the height-associated SNPs across the three continental populations from 1000 Genomes. The orange line represents the mean F_ST_ of the height-associated SNPs. The histogram represents the distribution of mean F_ST_ values of the sets of control SNPs.

**Figure 3.**
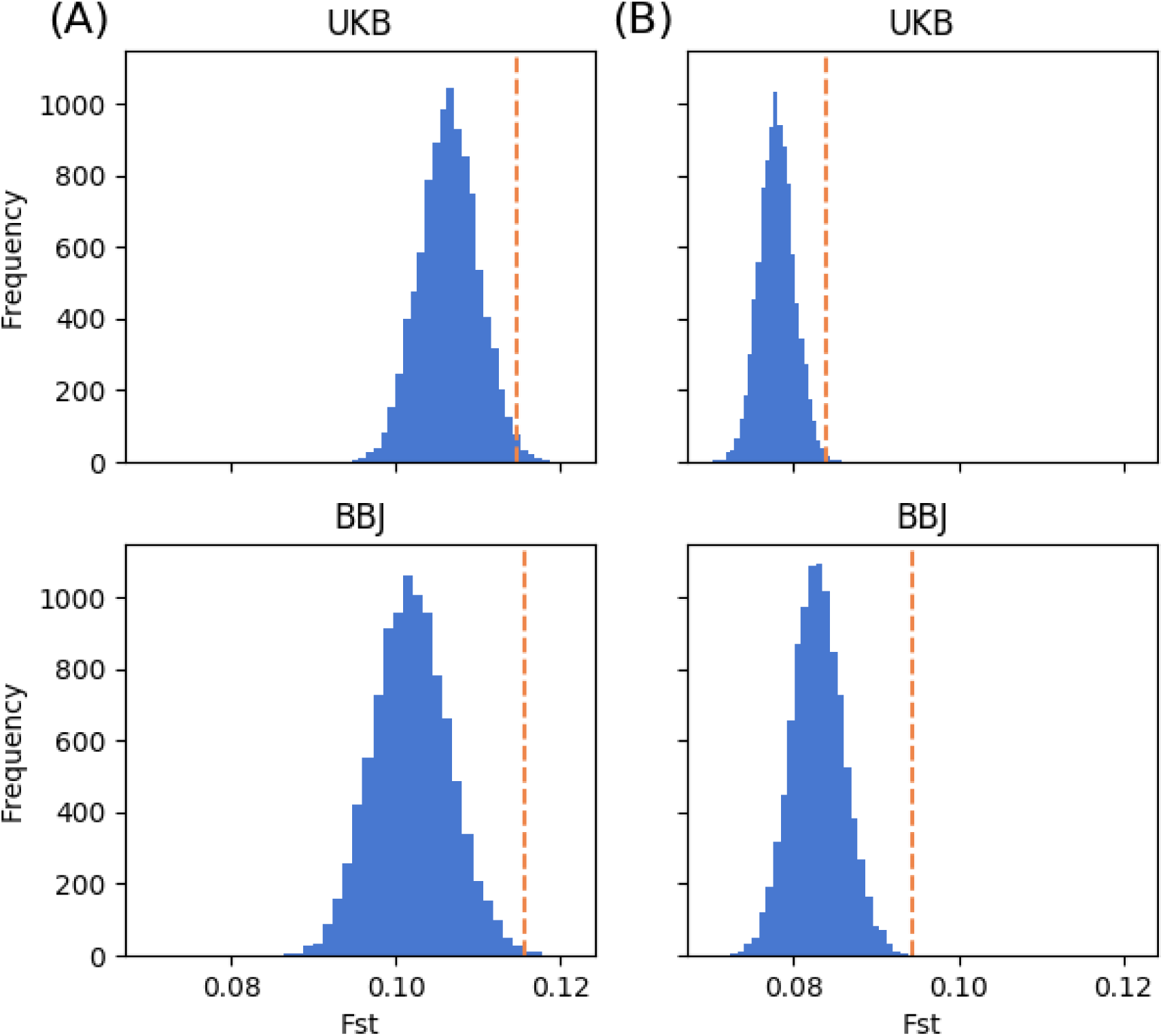
Mean F_ST_ values of the height-associated SNPs across the three continental populations (A) and across the seven regional populations (B) from HGDP. The orange line represents the mean F_ST_ of the height-associated SNPs. The histogram represents the distribution of mean F_ST_ values of the sets of control SNPs.

### PS-based analysis

On the basis of the height-associated SNPs ascertained from each of the three GWAS panels, we calculated the PSs for the three continental populations from 1000 Genomes. We used Berg and Coop’s Q_x_ and Conditional *Z* score framework to evaluate the significance of differences in PSs across populations and continents. We observed a clear signal of adaptation when we used height-associated SNPs ascertained from GIANT (*p* = 3.01e-6 for Q_x_ test). At the continental level, Europeans were significantly taller than would be expected under neutral drift given their genetic relationship with Africans and East Asians and the PSs in Africans and East Asians (*p* = 1.67e-14 for Conditional *Z* score); and East Asians were significantly shorter than neutral expectation (*p* = 3.11e-4 for Conditional *Z* score). However, the signals disappeared when we used height-associated SNPs ascertained from UKB (*p* = 0.278 for Q_x_ test) or BBJ (*p* = 0.426 for Q_x_ test) (**Figure 4**), suggesting the signal from Q_x_ analysis may be largely driven by uncorrected stratification in the GIANT data. Moreover, the ranked orders of populations based on PSs were also variable across GWAS panels, consistent with previous reports of poor transferability of PS models across continental populations (Martin *et al*. 2017; but see Ragsdale *et al*. 2020). For example, among the three continents, Europeans appeared to have the highest mean PS based on the height-associated SNPs ascertained from UKB, while Africans had the highest mean PS based on the SNPs ascertained from BBJ. Using Guo et al.’s PS-based approach, we also observed the adaptive signature in GIANT but not in UKB or BBJ (**Figure S9**).

**Figure 4.**
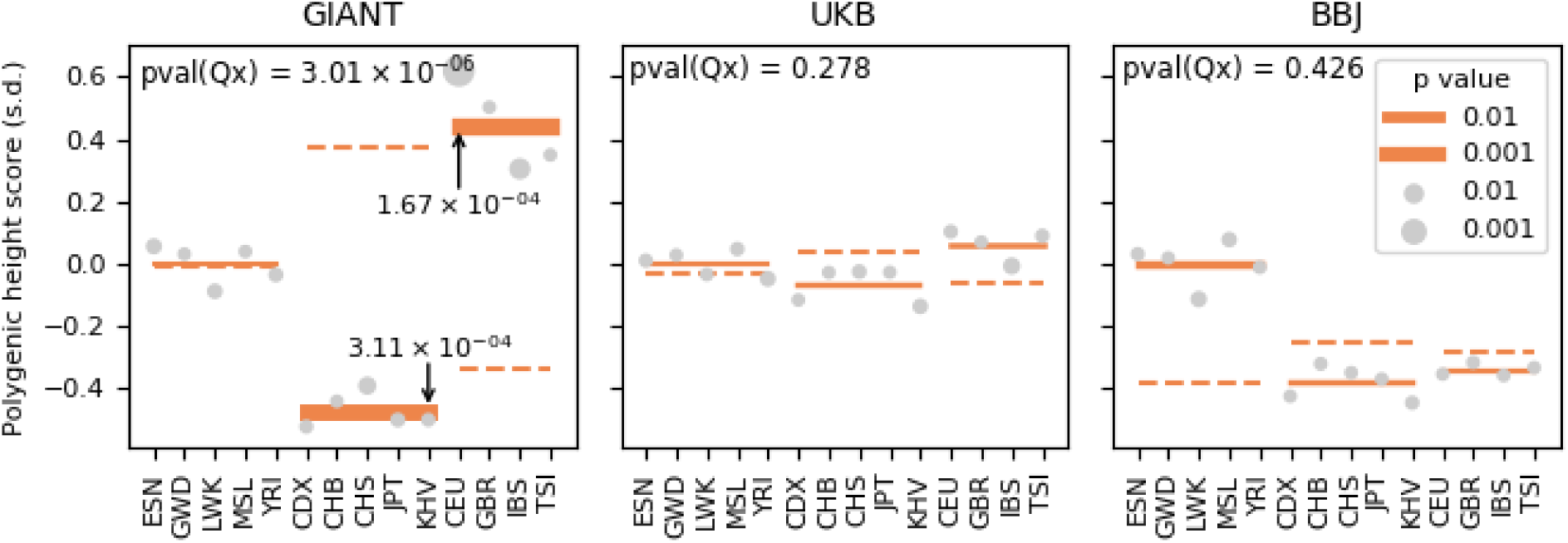
Q_x_ tests on PSs for the three continental populations from 1000 Genomes. The PSs were constructed on the basis of the height-associated SNPs ascertained from GIANT (A), UKB (B), and BBJ (C) GWAS summary statistics. Pval(Q_x_) denotes the *p* value for Q_x_ test. The *p* value for conditional *Z* score is represented by the size of each circle for each population, and two populations with *p* lower than 0.01 are CEU (*p* = 9.52e-5) and IBS (*p* = 0.0056) in the analysis using GIANT summary statistics; the *p* value is represented by the thickness of each solid line for each continent, and those lower than 0.01 are shown in the plot. The orange dashed line denotes the expected PS for each continent. African populations: ESN (Esan in Nigeria), GWD (Gambian in Western Divisions in the Gambia), LWK (Luhya in Webuye, Kenya), MSL (Mende in Sierra Leone), and YRI (Yoruba in Ibadan, Nigeria); East Asian populations: CDX (Chinese Dai in Xishuangbanna, China), CHB (Han Chinese in Beijing, China), CHS (Southern Han Chinese), JPT (Japanese in Tokyo, Japan), and KHV (Kinh in Ho Chi Minh City, Vietnam); European populations: CEU (Utah residents with ancestry from northern and western Europe), GBR (British in England and Scotland), IBS (Iberian Population in Spain), and TSI (Toscani in Italia).

We repeated the Q_x_ and Conditional *Z* score analysis using height-associated SNPs ascertained from UKB and BBJ in the three continental populations and the seven regional populations from HGDP and found no signature of selection at the continent level (**Figure S10 and S11**), recapitulating our observation from 1000 Genomes. In these analyses, the only consistently discernable signal of selection are observed in Sardinians, confirming our previous result that shorter height among Sardinians compared to other European populations may be a result of natural selection (Chen *et al*. 2020). Taken together, PS-based analysis in this context appears to be particularly susceptible to the choice of SNPs used in the analysis. While we observed a robust signature of differentiation using F_ST_ among continental populations, the adaptive signatures based on PS calculated from GIANT-ascertained SNPs disappeared when we used UKB- or BBJ-ascertained SNPs.

## Discussion

By ascertaining height-associated alleles from the UKB and BBJ GWAS panels, we showed that frequencies of these alleles were not impacted by population structure represented by the first two PCs of PCA across three continents (*i*.*e*. Africa, Europe, and East Asia, **Figure 1**). Using these two sets of height-associated SNPs, our study has confirmed that height alleles appear to be more differentiated among the three continental populations, compared to matched SNPs. Our F_ST_ results are qualitatively consistent with previous reports based summary statistics from GIANT (Guo *et al*. 2018) (**Figures 2 and 3**) and support a global signature of polygenic adaptation at height-associated SNPs. However, we could not detect a signature of polygenic adaptation using PS-based methods (**Figures 4, S9, S10, and S11**).

Although the signal of adaptation at height-associated SNPs is robust when using F_ST_ method, it is still not clear which population is under selection and contributes to the signal, since F_ST_ is not able to show the direction of differentiation. PS-based methods applied to multiple populations were expected to help address this question, but we could not identify significant differentiation in PS among the three continental populations, whether we used the Q_x_ test or Guo et al.’s PS framework (**Figures 4, S9, S10, and S11**). In fact, the ranked orders of populations based on PS of height-associated SNPs were sensitive to the ascertainment GWAS panel. Therefore, PS-based methods in the context of global populations appear to be particularly susceptible to the choice of SNPs used in the analysis and could not provide a robust signal indicating the populations under selection.

The robust signal in F_ST_-based but a lack of signal in PS-based analysis could be attributed to at least two possible reasons. The first reason could be the poor transferability of PS across populations. While polygenic scores for human complex traits have been shown to be significantly correlated with the observed phenotype within population (Martin *et al*. 2017, 2019; Khera *et al*. 2018; Mars *et al*. 2020), because of factors such as the divergence of causal allele frequency and differences in LD between populations, PS across continental populations are much less efficacious in predicting phenotypes (Martin *et al*. 2017, 2019; Dikilitas *et al*. 2020; Marnetto *et al*. 2020). Poorly correlated scores would severely impact the power and interpretation of natural selection inferences based on these scores. Therefore, it may be a good practice for inference of polygenic adaptation based on PS to first demonstrate that the scores used are sufficiently predictive of the phenotype being studied.

In principle, F_ST_-based inference of polygenic adaptation could also be affected by the same factors leading to poor transferability of PS, even though it does not take into account the estimated effect sizes or the direction of effects like the PS-based methods. The second reason could thus be the genetic redundancy of polygenic traits. That is, a single phenotype corresponds to a larger number of causal loci. The consequence of this feature is that independently adaptive populations could converge to the same trait optimum using different sets of alleles (Barghi *et al*. 2020). As a result, it could lead to a loss of power of PS-based methods to detect polygenic adaptation if more than one population are under selection. To illustrate this potential, we performed forward simulations and compared the power of Q_x_ and F_ST_ to detect signatures of polygenic adaptation (**Figure 5**). We simulated three scenarios of polygenic selection on 120 alleles but with one, two, or three populations under selection, each using a different set of alleles to evolve (**Methods**). In each case, the genetic architecture is perfectly known, removing the concerns of poor transferability of PSs across populations. When only one population is under selection, Q_x_ test generally had more power than F_ST_ across the spectrum of selective coefficients tested here, as previously demonstrated (Berg and Coop 2014) (**Figure 5A**). When two populations are under selection, using different sets of alleles, Q_x_ test also had more power than F_ST_ to identify that selection is happening (**Figure 5B**), although the conditional *Z* statistics would often indicate that the wrong population is under selection (**Figure S12**). Assuming the PSs are predictive of the phenotype in each population, in this case a follow-up within-population analysis, such as one using trajectory of polygenic scores (Edge and Coop 2019), could complement the Qx statistics and clarify the population under selective pressure (Chen *et al*. 2020). Finally, when all three populations are under selection, we saw that across the range of selection coefficient (from *s* = 5e-3 to 6e-4), F_ST_ attained greater power compared to Q_x_ (**Figure 5C**). Therefore, in a scenario with independent selections among multiple populations on the same genetic architecture of a complex trait, Qx statistics could lose power to infer the presence of adaptation while F_ST_ remains more robust. This seems to be a plausible explanation for our observations here for human height, as the trait architecture is likely largely shared among all humans (Kang *et al*. 2010; N’Diaye *et al*. 2011; Brick *et al*. 2019; Akiyama *et al*. 2019) but recent adaptations could occur independently in multiple populations; we know of at least two instances of adaptive signatures at height-associated SNPs, in Sardinia (Zoledziewska *et al*. 2015; Chen *et al*. 2020) and in Flores (Tucci *et al*. 2018), with other possible examples (Edge and Coop 2019; Berg *et al*. 2019; Sohail *et al*. 2019; Chen *et al*. 2020).

**Figure 5.**
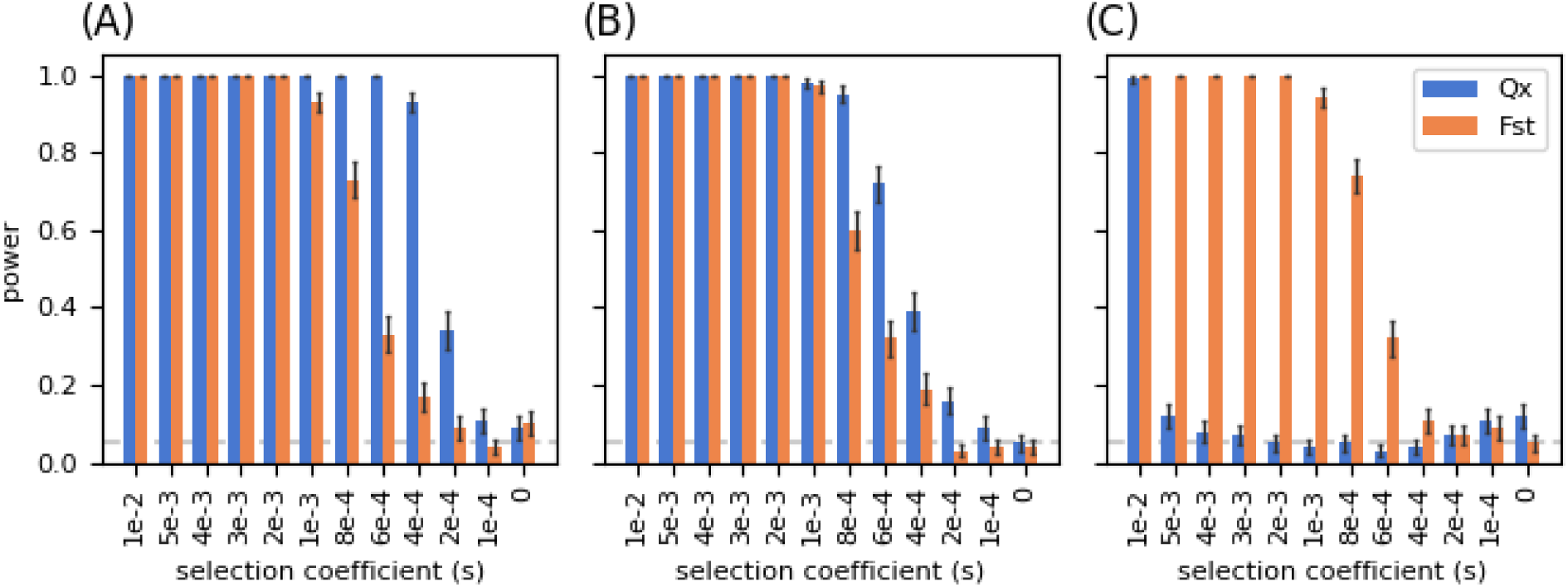
Forward simulation of power of Q_x_ and F_ST_ under different scenarios. (A) one population under selection; (B) two populations under selection; (C) three populations under selection.

Our study is motivated by the recent observations (Berg *et al*. 2019; Sohail *et al*. 2019) that residual population stratification in GWAS summary statistics in GIANT may have over-estimated the signal of polygenic adaptation at height-associated SNPs in the past. The impact of global population stratification on the effect size estimates in the GIANT summary statistics (**Figure 1**) could stem from shared ancestries between Europeans and other populations. The history of humans is a history of migration and admixture, and human populations are often connected by continuous gene flow (Hellenthal *et al*. 2014; Peter *et al*. 2020). As a result, shared history or gene flow between structured populations could result in stratifications both within and between populations. For example, the migration of early farmers during the Early Neolithic from the Near East and the migration of Yamnaya pastoralists during the Late Neolithic from the Yamnaya steppe contributed to the population structure observed between present-day Northern and Southern Europeans (Haak *et al*. 2015). Shared Yamnaya ancestry across Eurasia could also induce a correlation of shared structure between European and Asian populations. There is also evidence of recent gene flow (∼ 10 generations ago) from North Africa leading to the south-to-north latitudinal gradient of North African ancestry in Europe (Botigué *et al*. 2013), which may facilitate the correlation in allele frequencies between the European and African continents. Given the interconnectedness among human populations, it seems that the potential of residual stratification in a consortium GWAS (from a single, seemingly homogenous, population, nonetheless) to confound analyses across multiple populations should not be ignored.

There are multitudes of reasons to include more diverse populations into GWAS. The potential for genetic redundancy of a trait under selection would support greater inclusions in genetic analysis, as geographically diverse populations would provide opportunities to probe into different aspects of the trait architecture. However, it is already difficult to control for stratification in a within-continent population such as European-ancestry individuals, and the problem of stratification will only be exacerbated when we examine rarer and rarer variants (Mathieson and McVean 2012; Zaidi and Mathieson 2020). Therefore, we should bear in mind the interconnectedness across human populations, carefully characterize our genetic findings, and thread carefully when interpreting signals of selection (Novembre and Barton 2018).

## Acknowledgements

We gratefully thank Jing Guo and Jian Yang for providing the script for PS analysis. This work is supported by start-up funds provided by the Center for Genetic Epidemiology at the Keck School of Medicine of the University of Southern California (USC) (to C.W.K.C.). Computation for this work is supported by USC’s Center for Advanced Research Computing (https://carc.usc.edu)

## Reference

1000 Genomes Project Consortium, A. Auton, L. D. Brooks, R. M. Durbin, E. P. Garrison, et al., 2015 A global reference for human genetic variation. Nature 526: 68–74. https://doi.org/10.1038/nature15393

Akiyama M., K. Ishigaki, S. Sakaue, Y. Momozawa, M. Horikoshi, et al., 2019 Characterizing rare and low-frequency height-associated variants in the Japanese population. Nat. Commun. 10: 4393. https://doi.org/10.1038/s41467-019-12276-5

Barghi N., J. Hermisson, and C. Schlötterer, 2020 Polygenic adaptation: a unifying framework to understand positive selection. Nat. Rev. Genet. 1–13. https://doi.org/10.1038/s41576-020-0250-z

Berg J. J., and G. Coop, 2014 A Population Genetic Signal of Polygenic Adaptation, (M. W. Feldman, Ed.). PLoS Genet. 10: e1004412. https://doi.org/10.1371/journal.pgen.1004412

Berg J. J., A. Harpak, N. Sinnott-Armstrong, A. M. Joergensen, H. Mostafavi, et al., 2019 Reduced signal for polygenic adaptation of height in UK Biobank. Elife 8. https://doi.org/10.7554/eLife.39725

Bergström A., S. A. McCarthy, R. Hui, M. A. Almarri, Q. Ayub, et al., 2020 Insights into human genetic variation and population history from 929 diverse genomes. Science (80-.). 367: eaay5012. https://doi.org/10.1126/science.aay5012

Botigué L. R., B. M. Henn, S. Gravel, B. K. Maples, C. R. Gignoux, et al., 2013 Gene flow from North Africa contributes to differential human genetic diversity in southern europe. Proc. Natl. Acad. Sci. U. S. A. 110: 11791–11796. https://doi.org/10.1073/pnas.1306223110

Brick L. A., M. C. Keller, V. S. Knopik, J. E. McGeary, and R. H. C. Palmer, 2019 Shared additive genetic variation for alcohol dependence among subjects of African and European ancestry. Addict. Biol. 24: 132–144. https://doi.org/10.1111/adb.12578

Chang C. C., C. C. Chow, L. C. Tellier, S. Vattikuti, S. M. Purcell, et al., 2015 Second-generation PLINK: rising to the challenge of larger and richer datasets. Gigascience 4: 7. https://doi.org/10.1186/s13742-015-0047-8

Chen M., C. Sidore, M. Akiyama, K. Ishigaki, Y. Kamatani, et al., 2020 Evidence of Polygenic Adaptation in Sardinia at Height-Associated Loci Ascertained from the Biobank Japan. Am. J. Hum. Genet. https://doi.org/10.1016/j.ajhg.2020.05.014

Dikilitas O., D. J. Schaid, M. L. Kosel, R. J. Carroll, C. G. Chute, et al., 2020 Predictive Utility of Polygenic Risk Scores for Coronary Heart Disease in Three Major Racial and Ethnic Groups. Am. J. Hum. Genet. 106: 707–716. https://doi.org/10.1016/j.ajhg.2020.04.002

Edge M. D., and G. Coop, 2019 Reconstructing the History of Polygenic Scores Using Coalescent Trees. Genetics 211: 235–262. https://doi.org/10.1534/genetics.118.301687

Field Y., E. A. Boyle, N. Telis, Z. Gao, K. J. Gaulton, et al., 2016 Detection of human adaptation during the past 2000 years. Science (80-.). 354: 760–764. https://doi.org/10.1126/SCIENCE.AAG0776

Guo J., Y. Wu, Z. Zhu, Z. Zheng, M. Trzaskowski, et al., 2018 Global genetic differentiation of complex traits shaped by natural selection in humans. Nat. Commun. 9: 1865. https://doi.org/10.1038/s41467-018-04191-y

Haak W., I. Lazaridis, N. Patterson, N. Rohland, S. Mallick, et al., 2015 Massive migration from the steppe was a source for Indo-European languages in Europe. Nature 522: 207–211. https://doi.org/10.1038/nature14317

Haller B. C., and P. W. Messer, 2019 SLiM 3: Forward Genetic Simulations Beyond the Wright-Fisher Model, (R. Hernandez, Ed.). Mol. Biol. Evol. 36: 632–637. https://doi.org/10.1093/molbev/msy228

Hellenthal G., G. B. J. Busby, G. Band, J. F. Wilson, C. Capelli, et al., 2014 A Genetic Atlas of Human Admixture History. Science (80-.). 343: 747–751. https://doi.org/10.1126/science.1243518

Hirata M., Y. Kamatani, A. Nagai, Y. Kiyohara, T. Ninomiya, et al., 2017 Cross-sectional analysis of BioBank Japan clinical data: A large cohort of 200,000 patients with 47 common diseases. J. Epidemiol. 27: S9–S21. https://doi.org/10.1016/j.je.2016.12.003

Kang S. J., C. W. K. Chiang, C. D. Palmer, B. O. Tayo, G. Lettre, et al., 2010 Genome-wide association of anthropometric traits in African- and African-derived populations. Hum. Mol. Genet. 19: 2725–2738. https://doi.org/10.1093/hmg/ddq154

Khera A. V, M. Chaffin, K. G. Aragam, M. E. Haas, C. Roselli, et al., 2018 Genome-wide polygenic scores for common diseases identify individuals with risk equivalent to monogenic mutations. Nat. Genet. 50: 1219–1224. https://doi.org/10.1038/s41588-018-0183-z

Locke A. E., K. M. Steinberg, C. W. K. Chiang, S. K. Service, A. S. Havulinna, et al., 2019 Exome sequencing of Finnish isolates enhances rare-variant association power. Nature 1–6. https://doi.org/10.1038/s41586-019-1457-z

Loh P.-R., G. Tucker, B. K. Bulik-Sullivan, B. J. Vilhjálmsson, H. K. Finucane, et al., 2015 Efficient Bayesian mixed-model analysis increases association power in large cohorts. Nat. Genet. 47: 284–290. https://doi.org/10.1038/ng.3190

Marnetto D., K. Pärna, K. Läll, L. Molinaro, F. Montinaro, et al., 2020 Ancestry deconvolution and partial polygenic score can improve susceptibility predictions in recently admixed individuals. Nat. Commun. 11: 1–9. https://doi.org/10.1038/s41467-020-15464-w

Mars N., J. T. Koskela, P. Ripatti, T. T. J. Kiiskinen, A. S. Havulinna, et al., 2020 Polygenic and clinical risk scores and their impact on age at onset and prediction of cardiometabolic diseases and common cancers. Nat. Med. 26: 549–557. https://doi.org/10.1038/s41591-020-0800-0

Martin A. R., C. R. Gignoux, R. K. Walters, G. L. Wojcik, B. M. Neale, et al., 2017 Human Demographic History Impacts Genetic Risk Prediction across Diverse Populations. Am. J. Hum. Genet. 100: 635–649. https://doi.org/10.1016/J.AJHG.2017.03.004

Martin A. R., M. Kanai, Y. Kamatani, Y. Okada, B. M. Neale, et al., 2019 Clinical use of current polygenic risk scores may exacerbate health disparities. Nat. Genet. 51: 584–591. https://doi.org/10.1038/s41588-019-0379-x

Mathieson I., and G. McVean, 2012 Differential confounding of rare and common variants in spatially structured populations. Nat. Genet. 44: 243–246. https://doi.org/10.1038/ng.1074

N’Diaye A., G. K. Chen, C. D. Palmer, B. Ge, B. Tayo, et al., 2011 Identification, Replication, and Fine-Mapping of Loci Associated with Adult Height in Individuals of African Ancestry, (P. M. Visscher, Ed.). PLoS Genet. 7: e1002298. https://doi.org/10.1371/journal.pgen.1002298

Nagai A., M. Hirata, Y. Kamatani, K. Muto, K. Matsuda, et al., 2017 Overview of the BioBank Japan Project: Study design and profile. https://doi.org/10.1016/j.je.2016.12.005

Novembre J., and N. H. Barton, 2018 Tread Lightly Interpreting Polygenic Tests of Selection. Genetics 208: 1351–1355. https://doi.org/10.1534/genetics.118.300786

Okada Y., Y. Momozawa, S. Sakaue, M. Kanai, K. Ishigaki, et al., 2018 Deep whole-genome sequencing reveals recent selection signatures linked to evolution and disease risk of Japanese. Nat. Commun. 9. https://doi.org/10.1038/s41467-018-03274-0

Peter B. M., D. Petkova, and J. Novembre, 2020 Genetic landscapes reveal how human genetic diversity aligns with geography. Mol. Biol. Evol. 37: 943–951. https://doi.org/10.1093/molbev/msz280

Price A. L., M. E. Weale, N. Patterson, S. R. Myers, A. C. Need, et al., 2008 Long-Range LD Can Confound Genome Scans in Admixed Populations. Am. J. Hum. Genet. 83: 132–135.

Pritchard J. K., J. K. Pickrell, and G. Coop, 2010 The Genetics of Human Adaptation: Hard Sweeps, Soft Sweeps, and Polygenic Adaptation. Curr. Biol. 20: R208–R215.

Ragsdale A. P., D. Nelson, S. Gravel, and J. Kelleher, 2020 Lessons Learned from Bugs in Models of Human History. Am. J. Hum. Genet. 107: 583–588. https://doi.org/10.1016/j.ajhg.2020.08.017

Robinson M. R., G. Hemani, C. Medina-Gomez, M. Mezzavilla, T. Esko, et al., 2015 Population genetic differentiation of height and body mass index across Europe. Nat. Genet. Vol. 47: 34. https://doi.org/10.1038/ng.3401

Sohail M., R. M. Maier, A. Ganna, A. Bloemendal, A. R. Martin, et al., 2019 Polygenic adaptation on height is overestimated due to uncorrected stratification in genome-wide association studies. Elife 8. https://doi.org/10.7554/eLife.39702

Tucci S., S. H. Vohr, R. C. McCoy, B. Vernot, M. R. Robinson, et al., 2018 Evolutionary history and adaptation of a human pygmy population of Flores Island, Indonesia. Science 361: 511–516. https://doi.org/10.1126/science.aar8486

Turchin M. C., C. W. Chiang, C. D. Palmer, S. Sankararaman, D. Reich, et al., 2012 Evidence of widespread selection on standing variation in Europe at height-associated SNPs. Nat. Genet. 44: 1015–1019. https://doi.org/10.1038/ng.2368

Wang S. R., V. Agarwala, J. Flannick, C. W. K. Chiang, D. Altshuler, et al., 2014 Simulation of finnish population history, guided by empirical genetic data, to assess power of rare-variant tests in Finland. Am. J. Hum. Genet. 94: 710–720. https://doi.org/10.1016/j.ajhg.2014.03.019

Wood A. R., T. Esko, J. Yang, S. Vedantam, T. H. Pers, et al., 2014 Defining the role of common variation in the genomic and biological architecture of adult human height. Nat. Genet. 46: 1173–1186. https://doi.org/10.1038/ng.3097

Zaidi A. A., and I. Mathieson, 2020 Demographic history impacts stratification in polygenic scores. bioRxiv 2020.07.20.212530. https://doi.org/10.1101/2020.07.20.212530

Zoledziewska M., C. Sidore, C. W. K. Chiang, S. Sanna, A. Mulas, et al., 2015 Height-reducing variants and selection for short stature in Sardinia. Nat. Genet. 47: 1352–1356. https://doi.org/10.1038/ng.3403

