## Supplemental Material for "Allele frequency differentiation at height-associated SNPs among continental human populations"

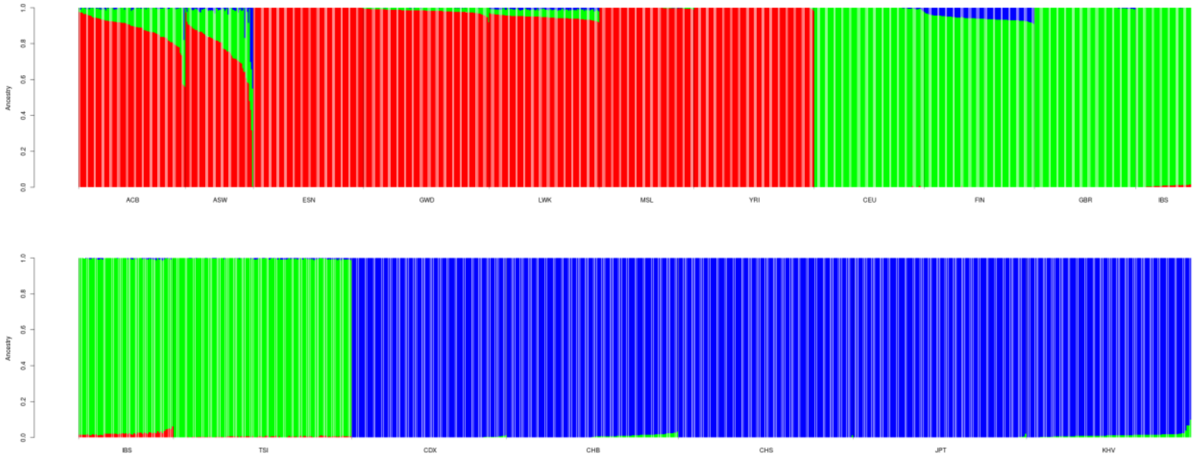


**Figure S1.** Ancestry composition of the three continental populations (Africans, Europeans, and East Asians) from 1000 Genomes using three ancestral populations ($K$ = 3) in ADMIXTURE analysis. The analysis was conducted using ADMIXTURE version 1.3.0 (Alexander *et al.* 2009) in unsupervised mode.

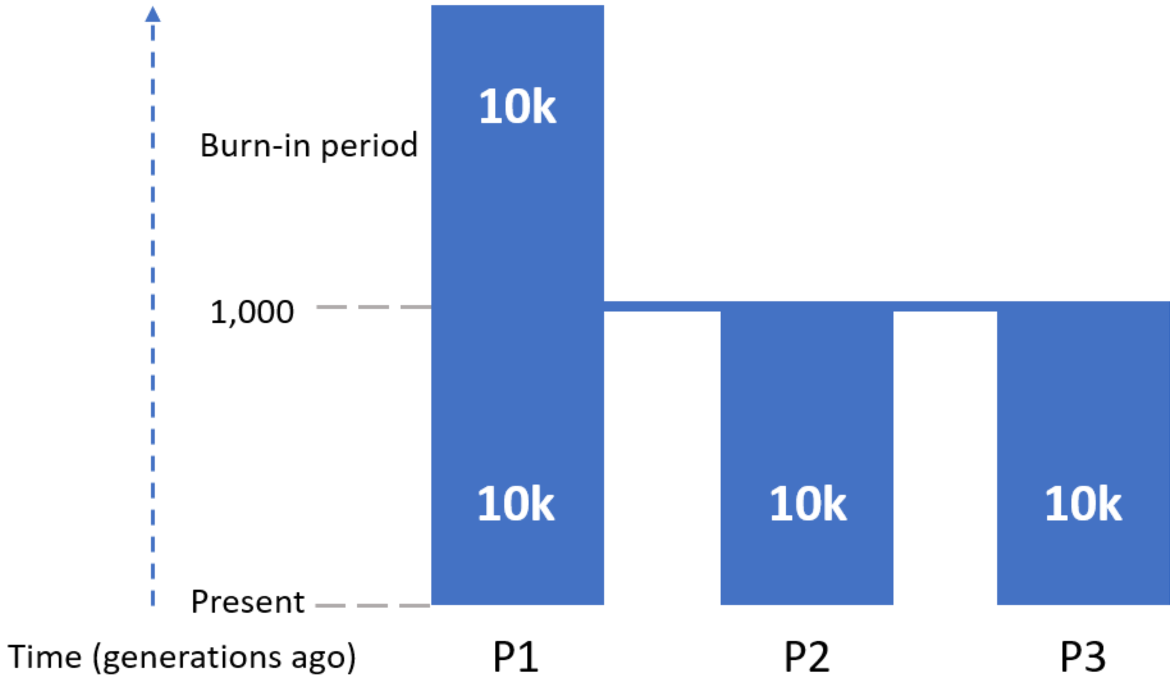


**Figure S2.** The demographic model used for the forward simulations. After a burn-in period of 10*Ne (100,000 generations), the ancestral population diverged into three subpopulations, P1, P2, and P3. In the presence of polygenic adaptation, immediately after the split, 120 randomly chosen independent mutations with global MAF > 20% were chosen to be either beneficial or deleterious in one of the three populations depending on the simulation scenario as described in the Method section. The simulation was run for 1,000 additional generations.

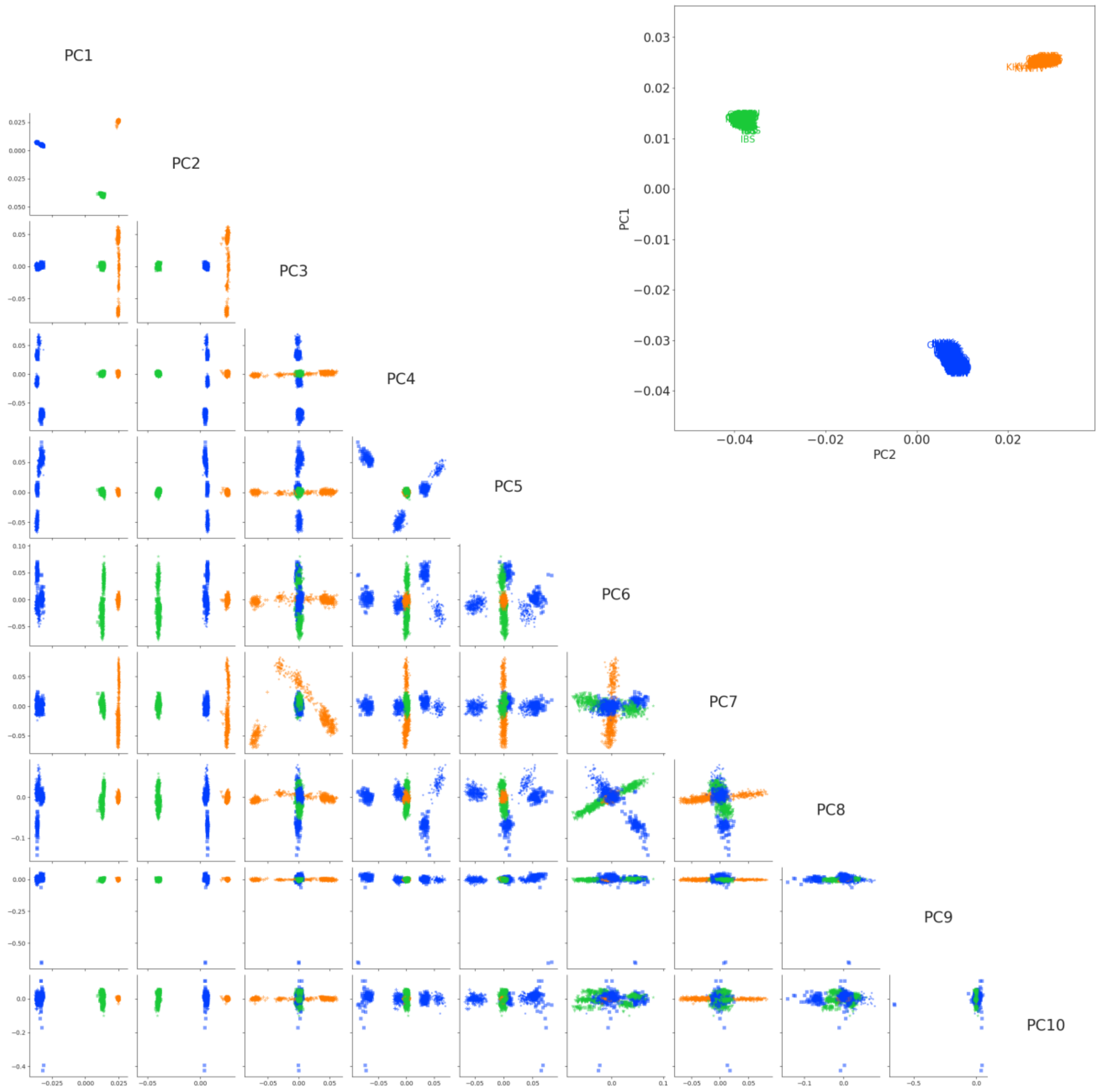


**Figure S3**. First ten principal components from PCA of the three continental populations from 1000 Genomes. Inset shows the first two principal components in greater detail. Colors in the triangular plot reflect that shown in the inset.

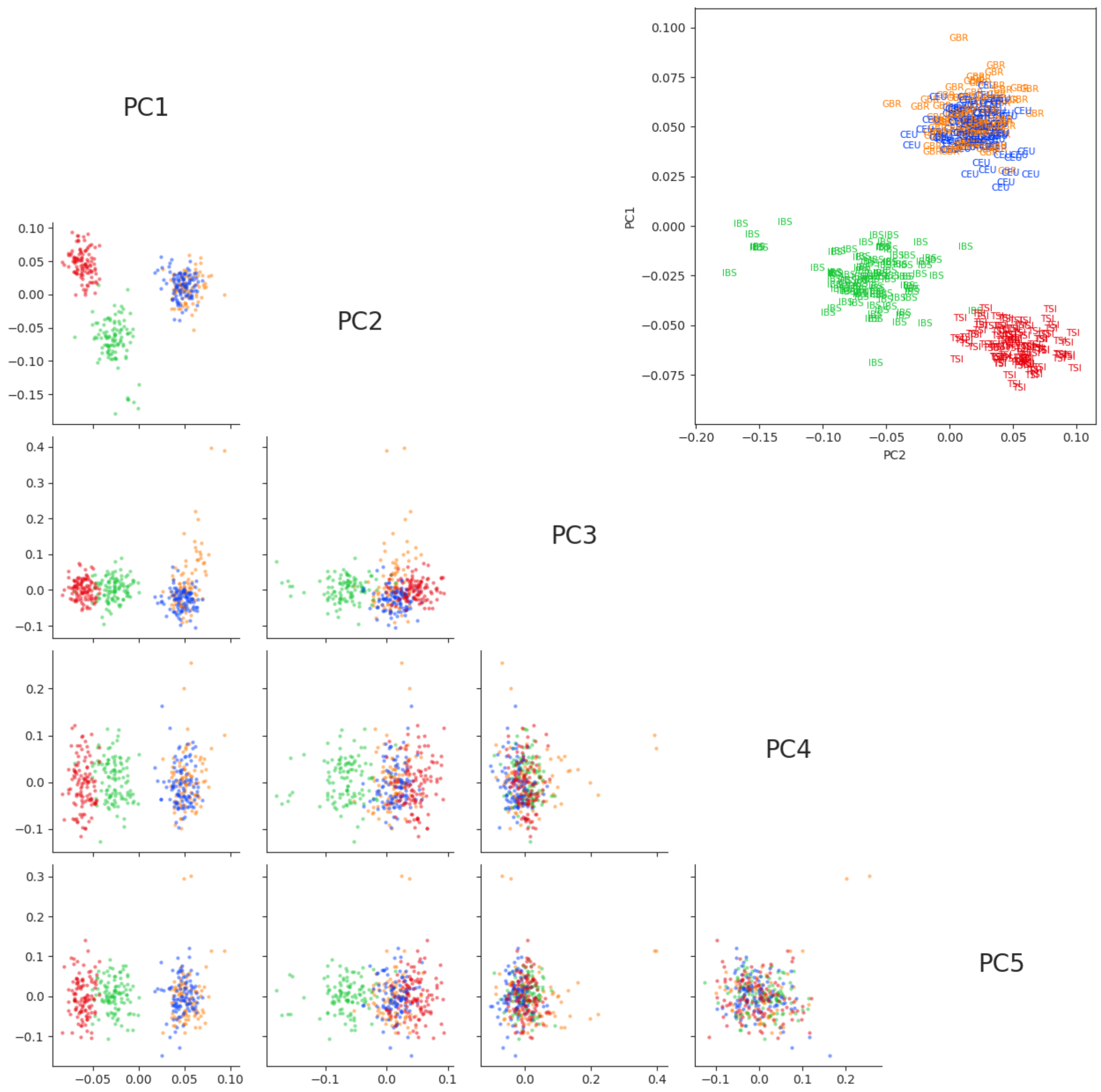


**Figure S4.** First five principal components from PCA of the four European populations from 1000 Genomes. Inset shows the first two principal components in greater detail. Colors in the triangular plot reflect that shown in the inset.

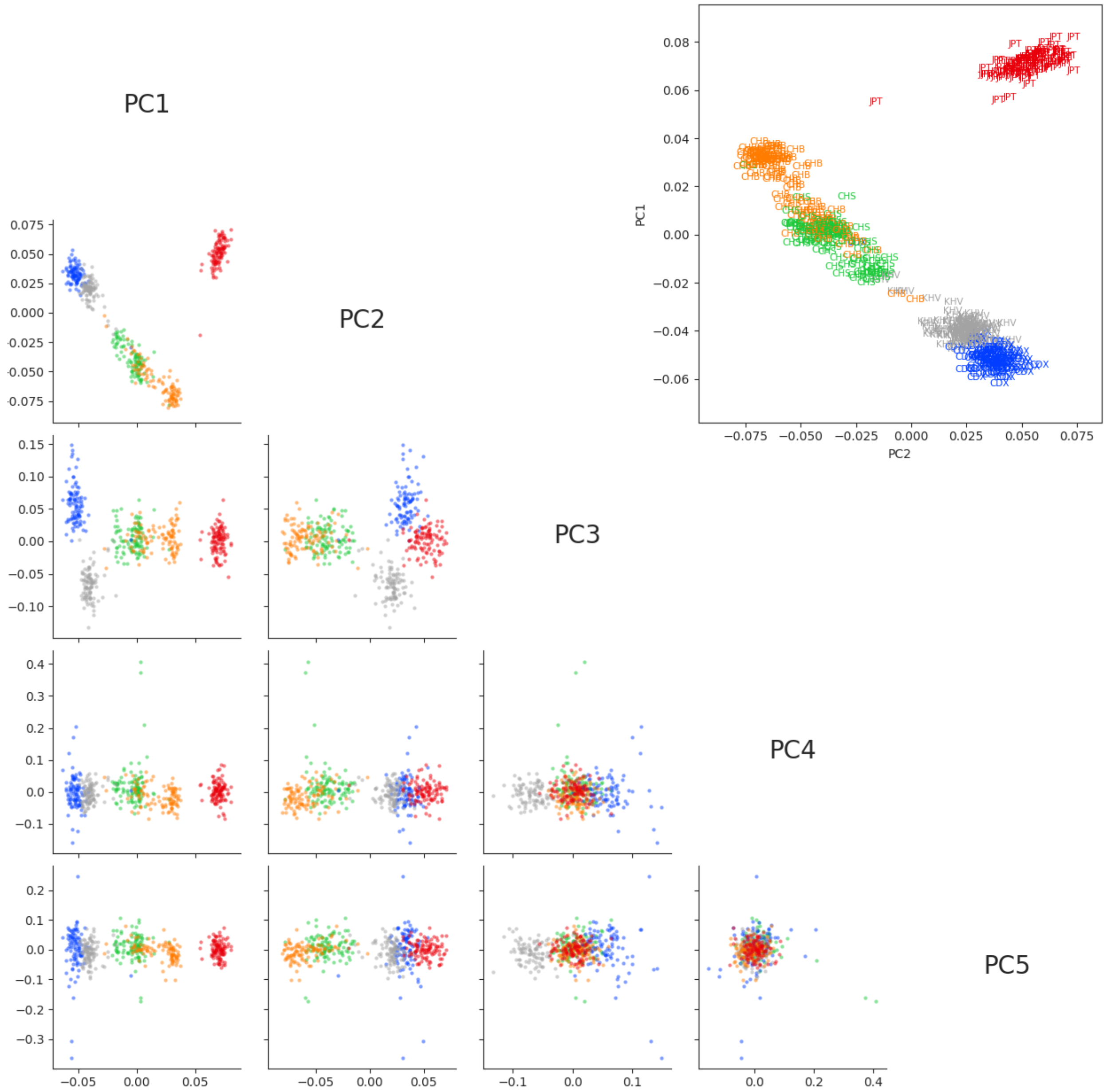


**Figure S5.** First five principal components from PCA of the five East Asian populations from 1000 Genomes. Inset shows the first two principal components in greater detail. Colors in the triangular plot reflect that shown in the inset.

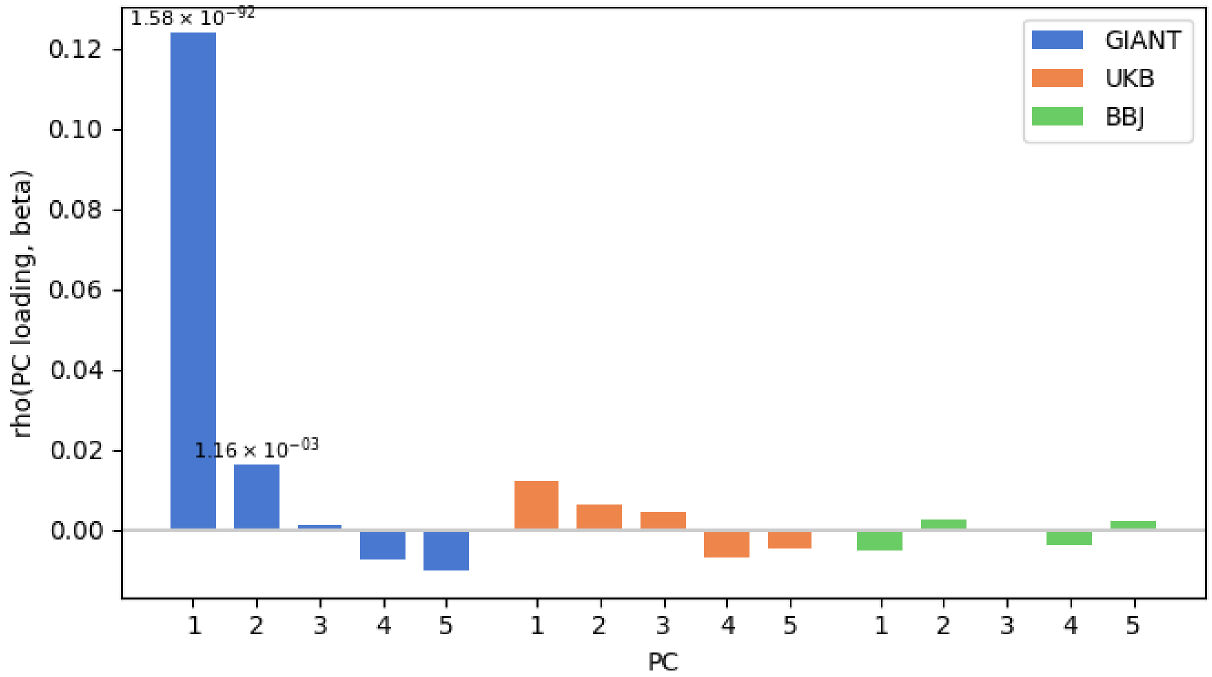


**Figure S6.** Evidence of stratification within Europe in GWAS summary statistics. Pearson correlation coefficients of PC loadings and SNP effects from GIANT, UKB, and BBJ based on all SNPs with MAF > 1% in each GWAS panel. Twenty PCs were computed in the four European populations from 1000 Genomes. *P* values are based on jackknife standard errors (1,000 blocks). *P* values lower than 0.05/20 are indicated on each bar. Only the fist 5 PCs are shown for readability.

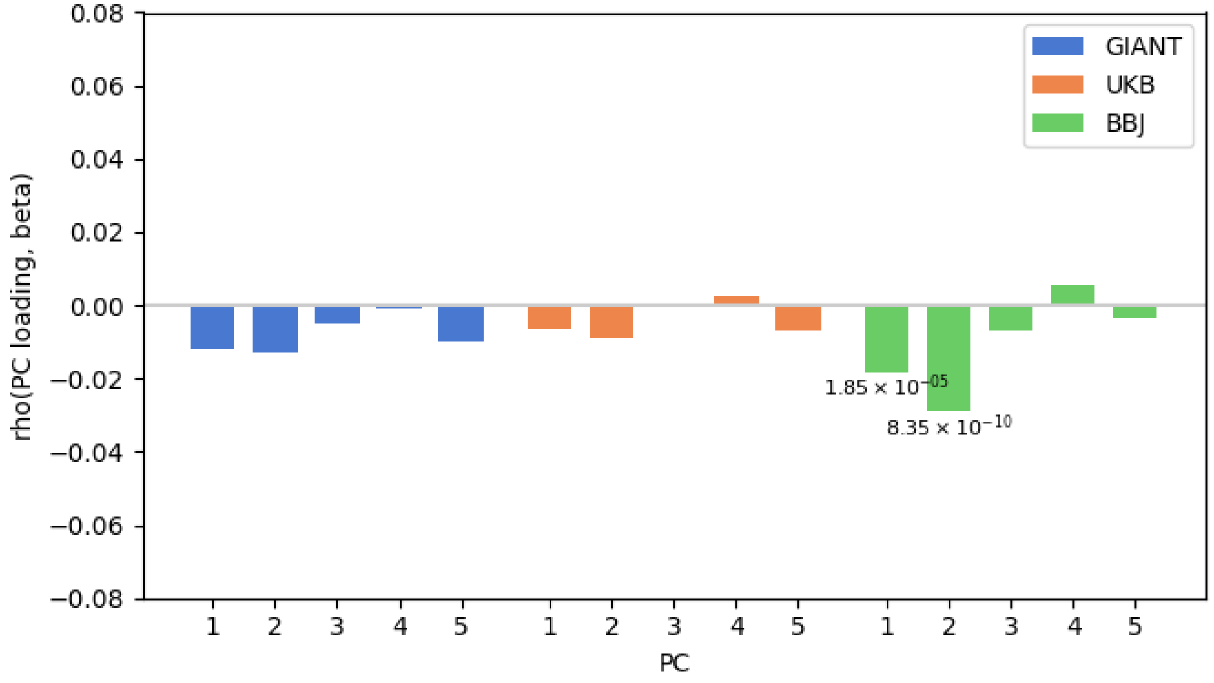


**Figure S7.** Evidence of stratification within East Asian in GWAS summary statistics. Pearson correlation coefficients of PC loadings and SNP effects from GIANT, UKB, and BBJ based on all SNPs with MAF > 1% in each GWAS panel. Twenty PCs were computed in the five East Asian populations from 1000 Genomes. *P* values are based on jackknife standard errors (1,000 blocks). *P* values lower than 0.05/20 are indicated on each bar. Only the first 5 PCs are shown for readability.

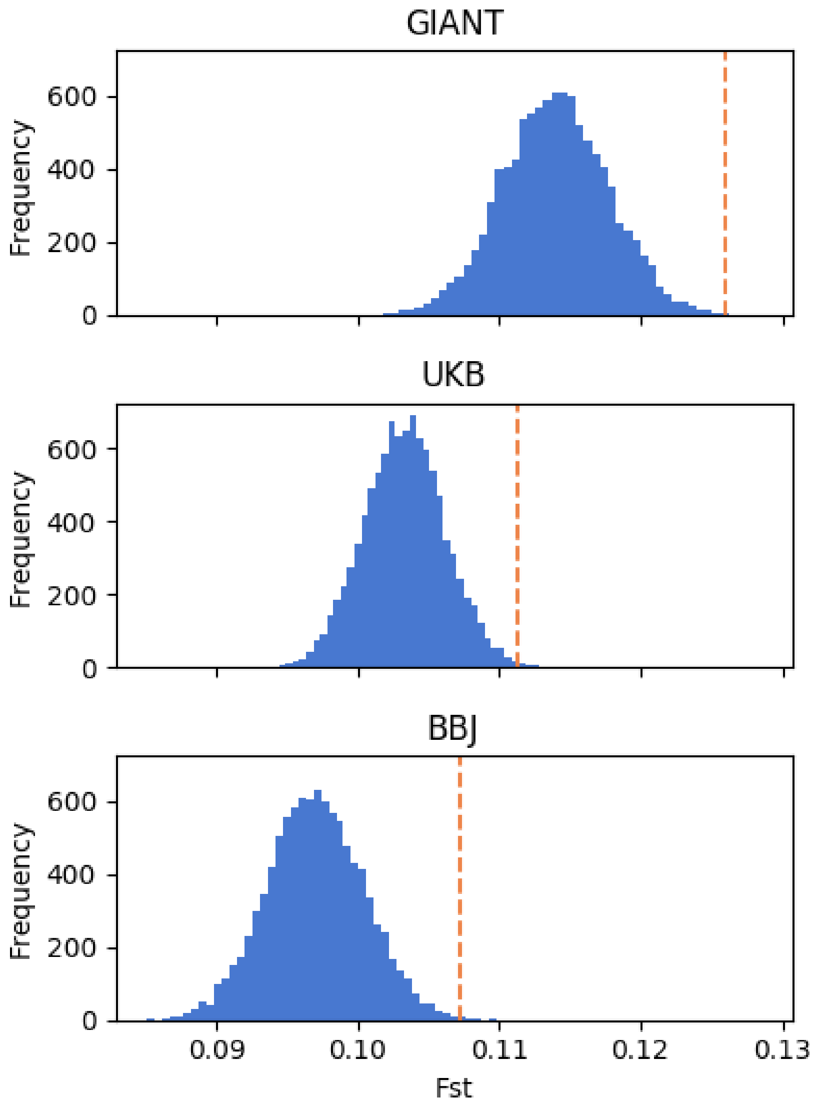


**Figure S8.** Mean $F_{ST}$ values of the height-associated SNPs with $p$ < 5e-6 across the three continental populations from 1000 Genomes. The orange line represents the mean $F_{ST}$ of the height-associated SNPs. The histogram represents the distribution of mean $F_{ST}$ values of the sets of control SNPs.

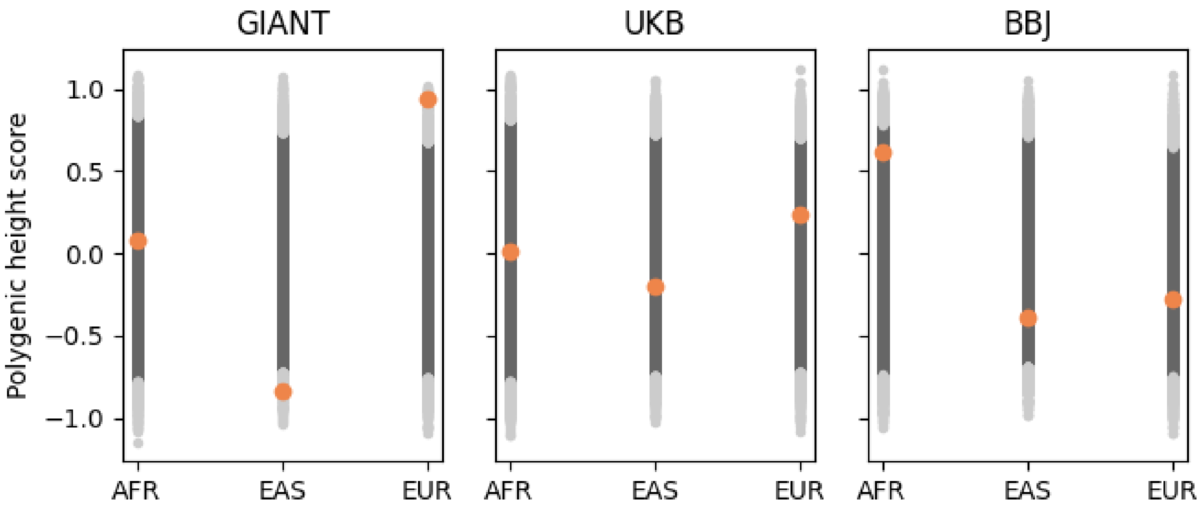


**Figure S9.** The mean deviation of the mean PS of a population from the overall mean. The blue dots represent the estimated deviation (in s.d. unites) of the mean PS based on the height-associated SNPs of a population from the overall mean across populations. The black dots represent the 95% confidence interval of the distribution of mean PS values of 10,000 sets of control SNPs; the gray dots represent the rest 5% mean PS values. AFR: Africans; EAS: East Asians; EUR: Europeans.

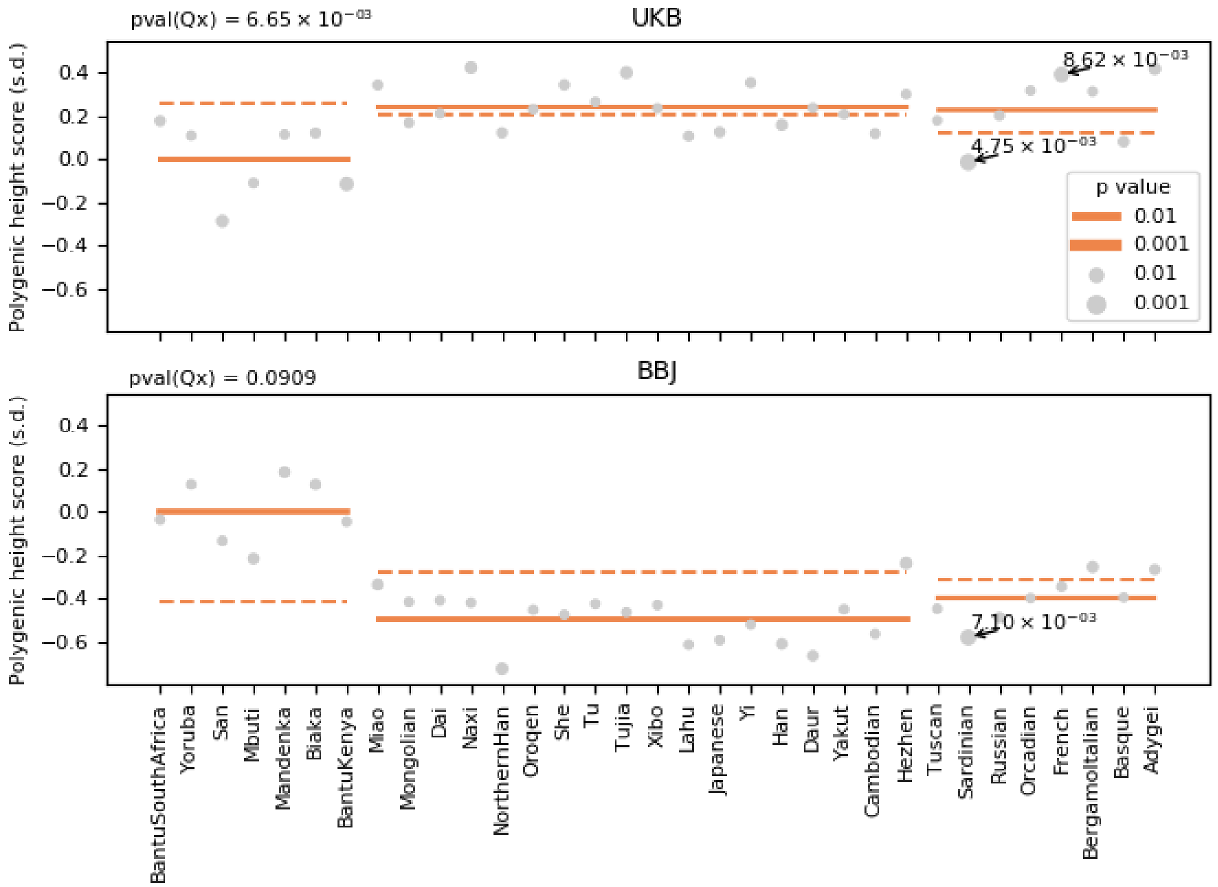


**Figure S10.** $Q_x$ tests on PS for the three continental populations from HGDP. The PS was constructed on the basis of the height-associated SNPs ascertained from UKB and BBJ GWAS summary statistics. Pval($Q_x$) denotes the *p* value for $Q_x$ test. The *p* value for conditional *Z* score is represented by the size of each circle for each population and by the thickness of each solid line for each continent, and those lower than 0.01 are shown in the plot. The orange dashed line denotes the expected PS for each continent.

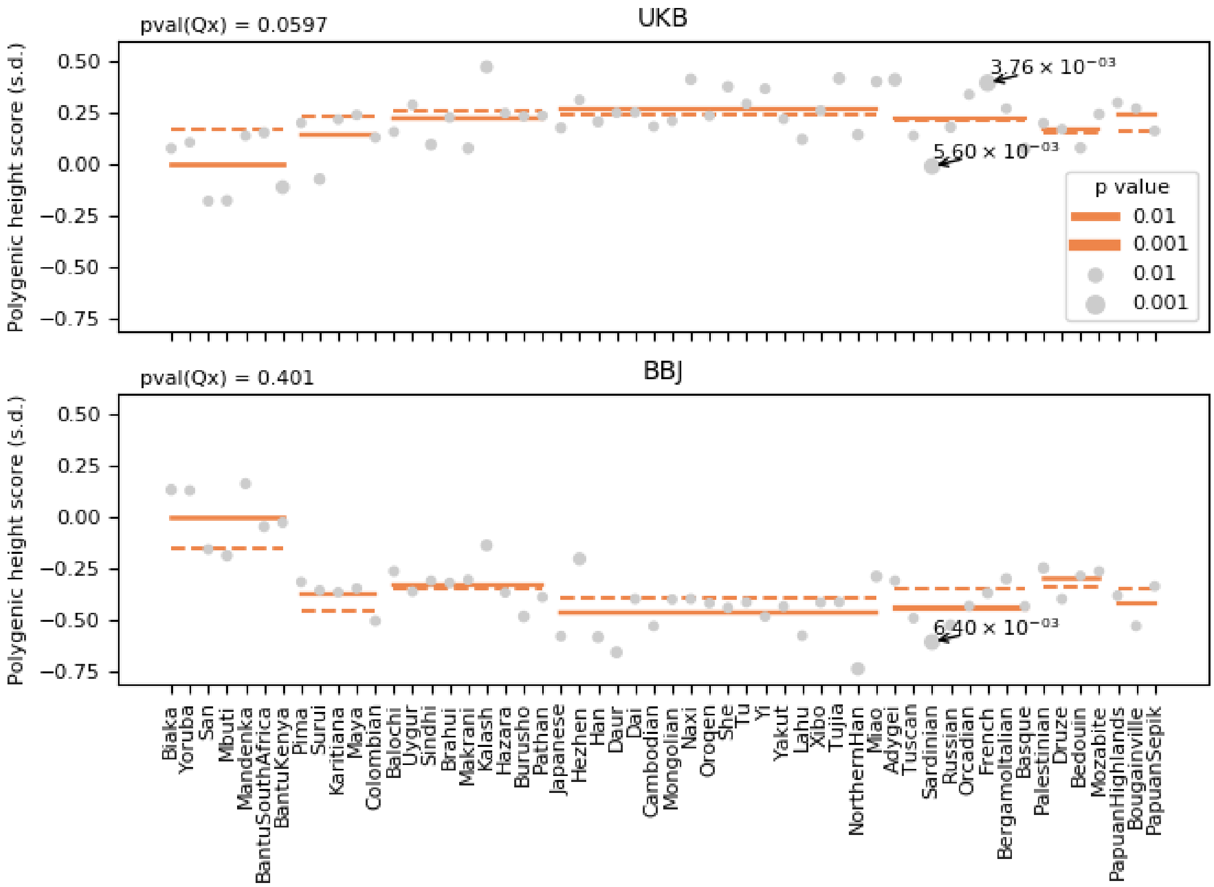


**Figure S11.** $Q_x$ tests on PS for the seven regional populations from HGDP. The PS was constructed on the basis of the height-associated SNPs ascertained from UKB and BBJ GWAS summary statistics. Pval($Q_x$) denotes the *p* value for $Q_x$ test. The *p* value for conditional *Z* score is represented by the size of each circle for each population and by the thickness of each solid line for each continent, and those lower than 0.01 are shown in the plot. The orange dashed line denotes the expected PS for each continent.

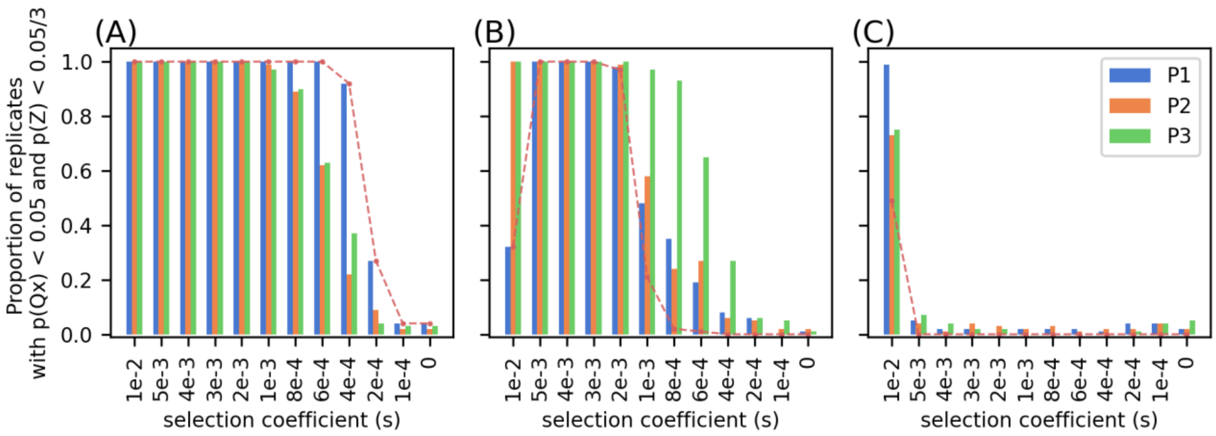


**Figure S12.** Proportion of replicates with significant $Q_x$ statistics ($p < 0.05$) and significant conditional $Z$ score ($p < 0.05/3$) for each population under different scenarios of forward simulation. For the conditional Z score, since three populations are tested, we used a Bonferroni-corrected threshold of 0.05/3. (A) one population (P1) under selection; (B) two populations (P1 and P2) under selection; (C) three populations (P1, P2, and P3) under selection. Red dash line indicates the proportion of replicates in which the population under selection is correctly identified. If more than one population is under selection (in (B) and (C)), all populations under selection need to be identified. In (B), even though P3 is not under selection, the conditional Z score often attributed the signature of selection to P3 instead of P1 and P2, particularly when the strength of selection is weak, around 1e-3 to 4e-4. A total of 100 replicates were performed in simulation.

**Supp Table 1**. Correlation between Fst across continents and absolute PC loading on PCA conducted across continents. We assessed if the correlation is significantly different from 0 by computing a *p* values based on jackknife standard errors (1,000 blocks). In all cases, the *p* values are hugely significant ($p < 1e\text{-}200$).

| | Global PCA (Figure S3) | |
|---|---|---|
| | PC1 | PC2 |
| GIANT | 0.632 | 0.199 |
| UKB | 0.467 | -0.124 |
| BBJ | 0.0983 | -0.132 |

### Reference

Alexander D. H., J. Novembre, and K. Lange, 2009 Fast model-based estimation of ancestry in unrelated individuals. Genome Res. 19: 1655–1664. https://doi.org/10.1101/gr.094052.109
